# *Sinorhizobium meliloti* BR-bodies promote fitness during host colonization

**DOI:** 10.1101/2024.04.05.588320

**Authors:** Kaveendya S. Mallikaarachchi, Jason L. Huang, Shanmukha Madras, Rodrigo A. Cuellar, Zhenzhong Huang, Alisa Gega, Imalka W. Rathnayaka-Mudiyanselage, Nadra Al-Husini, Natalie Saldaña-Rivera, Loi H. Ma, Eric Ng, Joseph C. Chen, Jared M. Schrader

**Author notes:** Corresponding Authors Jared M. Schrader, Joseph C. Chen.

## Abstract

Biomolecular condensates, such as the nucleoli or P-bodies, are non-membrane-bound assemblies of proteins and nucleic acids that facilitate specific cellular processes. Like eukaryotic P-bodies, the recently discovered bacterial ribonucleoprotein bodies (BR-bodies) organize the mRNA decay machinery, yet the similarities in molecular and cellular functions across species have been poorly explored. Here, we examine the functions of BR-bodies in the nitrogen-fixing endosymbiont *Sinorhizobium meliloti*, which colonizes the roots of compatible legume plants. Assembly of BR-bodies into visible foci in *S. meliloti* cells requires the C-terminal intrinsically disordered region (IDR) of RNase E, and foci fusion is readily observed *in vivo*, suggesting they are liquid-like condensates that form via mRNA sequestration. Using Rif-seq to measure mRNA lifetimes, we found a global slowdown in mRNA decay in a mutant deficient in BR-bodies, indicating that compartmentalization of the degradation machinery promotes efficient mRNA turnover. While BR-bodies are constitutively present during exponential growth, the abundance of BR-bodies increases upon cell stress, whereby they promote stress resistance. Finally, using *Medicago truncatula* as host, we show that BR-bodies enhance competitiveness during colonization and appear to be required for effective symbiosis, as mutants without BR-bodies failed to stimulate plant growth. These results suggest that BR-bodies provide a fitness advantage for bacteria during infection, perhaps by enabling better resistance against the host immune response.

**Significance:** While eukaryotes often organize their biochemical pathways in membrane-bound organelles, bacteria generally lack such subcellular structures. Instead, membraneless compartments called biomolecular condensates have recently been found in bacteria to enhance biochemical activities. Bacterial ribonucleoprotein bodies (BR-bodies), as one of the most widespread biomolecular condensates identified to date, assemble the mRNA decay machinery via the intrinsically disordered regions (IDRs) of proteins. However, the implications of such assemblies are unclear. Using a plant-associated symbiont, we show that the IDR of its mRNA degradation protein is necessary for condensate formation. Absence of BR-bodies results in slower mRNA decay and ineffective symbiosis, suggesting that BR-bodies play critical roles in regulating biochemical pathways and promoting fitness during host colonization.

## Introduction

Eukaryotic and bacterial cells have been found to compartmentalize their mRNA decay machinery into biomolecular condensates (1–3), non-membrane-bound organelles that organize critical cellular processes, including RNA degradation and storage. Bacterial ribonucleoprotein bodies (BR-bodies) were the first biomolecular condensate discovered in bacteria, composed of the RNA degradosome complex (4). The large intrinsically disordered region (IDR) of RNase E was found to be necessary and sufficient for BR-body formation, and bioinformatic analyses of other bacteria suggest that BR-bodies are likely widespread across phylogeny (2, 4–7), including for other RNases like RNase Y and J (8, 9). Initial characterization of BR-bodies was performed in *Caulobacter crescentus*, where it was shown that they assemble with poorly translated mRNAs and promote the rapid decay of mRNA (10, 11). While constitutively present, BR-bodies were also stimulated to assemble in various stress conditions, where they promoted resistance (4). While BR-bodies have been identified in other bacteria, including *Sinorhizobium meliloti* (4), the functional role of BR-bodies in mRNA decay and stress resistance has not been thoroughly tested across species. However, transposon insertions or targeted mutations in RNase E that truncate the large IDR, and likely abolish BR-body assembly, have been found to cause stress sensitivity, defects in host colonization, and other pleiotropic phenotypes (4, 12–18). This suggests that BR-bodies may promote similar molecular and cellular phenotypes across species.

*S. meliloti* is an α-proteobacterium that can live freely or in symbiosis with legume plants (19, 20). In order to colonize compatible hosts, *S. meliloti* must invade plant root tissue and survive host defenses, such as antimicrobial peptides (21–24). Following reciprocal signaling events between the bacterium and plant, *S. meliloti* establishes chronic infection by inducing formation of root nodules and colonizing them (25). After engulfment by plant cells in the nodules, changes in gene expression reprogram the bacteria for differentiation into nitrogen-fixing bacteroids, allowing the mutualistic infection to promote robust plant growth (19). In addition to transcriptional regulation of nitrogen fixation genes, post-transcriptional regulation by small RNAs have been found to impact a variety of cellular processes, including metabolism, the cell cycle, and quorum sensing (26, 27). RNase E in *S. meliloti* has been found to affect quorum sensing and S-adenosylmethionine homeostasis (28, 29), yet microarray analysis of wild type and a mini Tn5-insertion mutant identified only a small subset of mRNAs with altered steady-state levels when the IDR of RNase E was disrupted (29). However, measurements of steady-state mRNA cannot distinguish differences in the rates of mRNA turnover. Moreover, it has not been established whether the *S. meliloti* RNase E IDR is necessary for BR-body formation *in vivo*, or whether BR-bodies impact plant colonization and symbiosis.

To examine the conservation of molecular and cellular phenotypes affected by BR-bodies, we generated a C-terminal IDR truncation mutation in *S. meliloti* RNase E. We found that the IDR was necessary for BR-body formation and that the IDR deletion mutant showed a general slow-down in mRNA decay, suggesting that BR-bodies promote mRNA turnover. Through a combination of drug and stress treatments, we determined that *S. meliloti* BR-bodies are constitutively present, stimulated by poorly translated mRNA, and more strongly induced under a variety of stresses. Finally, we found that BR-bodies promote fitness during stress and *Medicago truncatula* root colonization. Altogether, these results suggest that many of the molecular and cellular activities promoted by BR-bodies are likely conserved across bacterial species, enhancing viability across diverse lifestyles.

## Results

### Sinorhizobium meliloti RNase E forms BR-bodies that promote mRNA decay

α-proteobacterial RNase E proteins contain an N-terminal catalytic domain and a large IDR in the C-terminal domain, which is necessary and sufficient for phase separation into BR-bodies in *C. crescentus* (Fig 1A, B) (2, 4). To generate a mutant of *S. meliloti* which cannot assemble BR-bodies, we truncated the IDR at codon 665 based upon the domain annotation (30) and predicted region of disorder from dSCOPE (31) (Fig 1B, C). This IDR has been defined previously to contain an alternating pattern of “charge patches” of positively and negatively charged amino acids which promote RNase E phase-separation through electrostatic interactions (Fig 1D) (2, 4). The IDR deletion mutanat (RNase EΔIDR) was viable and showed no detectable difference in growth rate compared to wild type in TY media (Fig S1). To examine the importance of the IDR in BR-body formation *in vivo*, the localization of RNase E-YFP and RNase EΔIDR-YFP was determined by fluorescence microscopy. RNase E-YFP showed robust formation of fluorescent foci, while RNase EΔIDR-YFP led to a near total loss of foci compared to the wild-type (Fig 2A). Consistent with previous observations in the related α-proteobacterium *C. crescentus* (4), this result suggests that the RNase EΔIDR mutation abolishes the ability to form BR-bodies *in vivo. C. crescentus* RNase E foci fusion events were observed via time-lapse microscopy, suggesting the foci are phase separated condensates *in vivo* (4). To test if *S. meliloti* RNase E can phase separate *in vivo*, we performed time-lapse microscopy with the RNase E-YFP strain (Fig 2B). Here, we similarly observed that *S. meliloti* RNase E-YFP foci could fuse like those of *C. crescentus*, suggesting they also form phase separated condensates *in vivo. C. crescentus* BR-bodies were shown to phase-separate with RNA both *in vivo* and *in vitro* (4, 11). A combination of genetic mutations and drug treatments suggested that *C. crescentus* RNase E competes with ribosomes *in vivo* for access to mRNAs which act as the major substrate for BR-body formation (4, 10). To test the impact of mRNA availability in *S. meliloti*, we performed growth assays in different conditions, including drug treatments with rifampicin, which blocks transcription and rapidly depletes the cell of mRNA, and chloramphenicol, which freezes ribosomes on mRNAs and traps mRNAs in polysomes (4)[Fig S2]. Upon short treatment of rifampicin (100 μg/mL for 30’) or chloramphenicol (200 μg/mL for 30’), we saw a strong reduction in the number of BR-bodies (Fig S2). This reduction suggests that *S. meliloti* BR-bodies assemble via the accumulation of poorly translated mRNAs with RNase E, as in *C. crescentus* (4).

**Figure 1:**
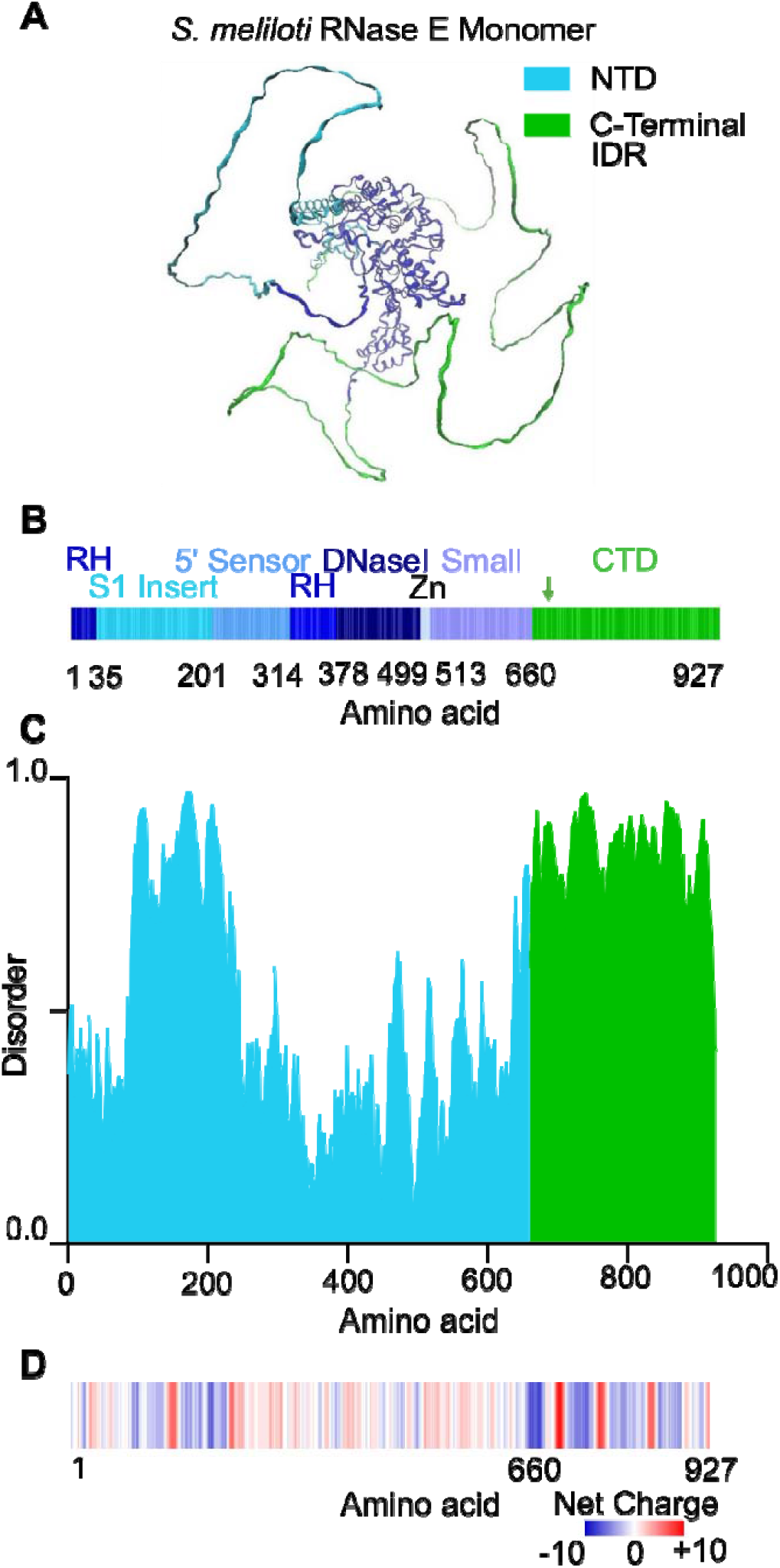
*S. meliloti* RNase E has the features necessary for phase separation and BR-body formation. **A.)** RNase E monomer structure predicted from AlphaFold (45, 46). The folded catalyti nuclease domain is colored in blue, while the IDR region is highlighted in green. **B.)** RNase E domain structure. Domain annotations were from Hardwick *et al*. (2011) (30). C.) Disorder across RNase E. dSCOPE predictions of the RNase E disorder (31). **D.)** Charge patterning across RNase E. The IDR i highly enriched with alternating charge patches, which were shown to be necessary and sufficient for phase separation in *C. crescentus* RNase E (4).

**Figure 2:**
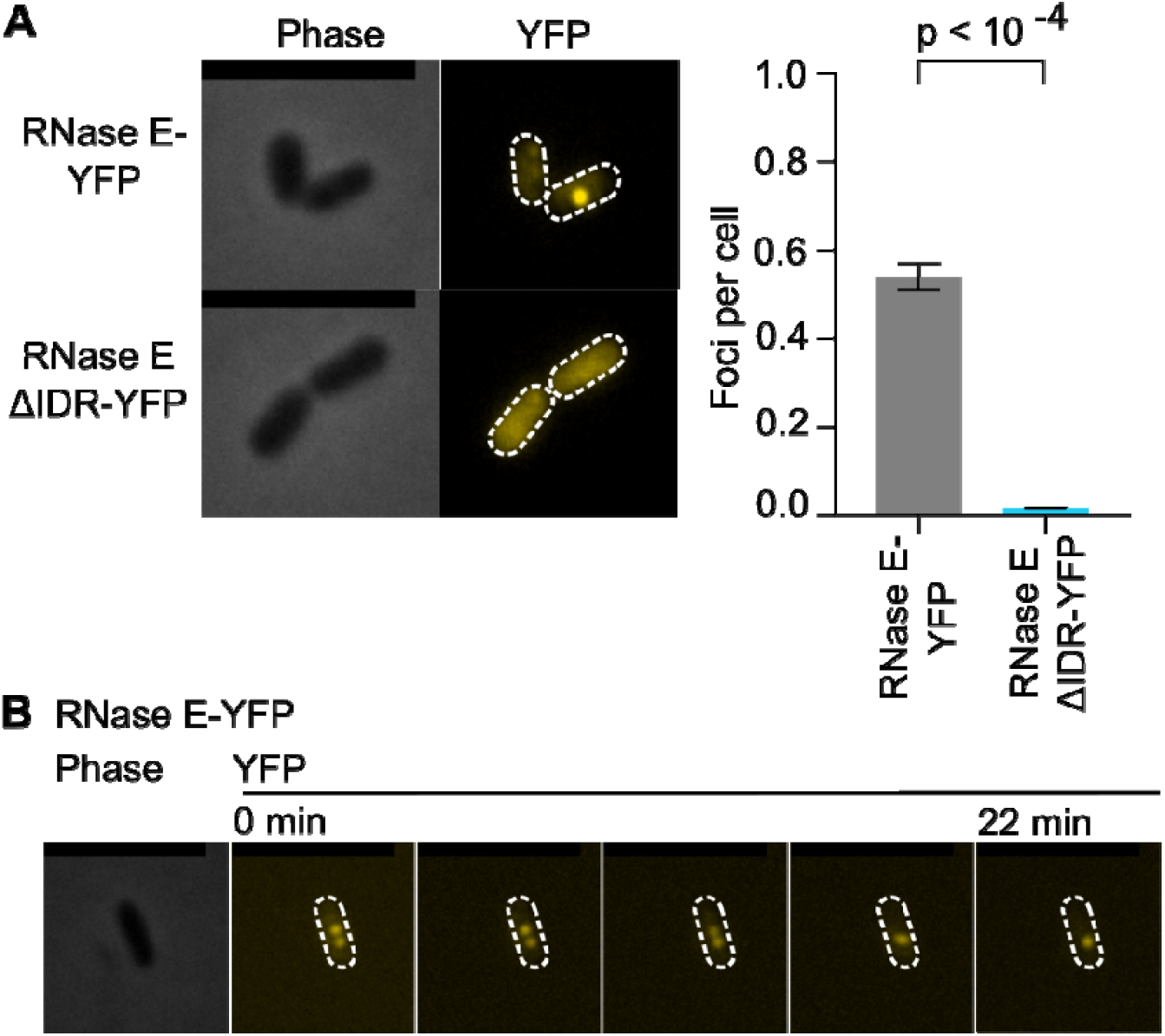
*S. meliloti* RNase E requires the C-terminal IDR to assemble into dynamic BR-bodies. **A.)** RNase E foci require the C-terminal IDR. Rm1021 cells harboring YFP fusion to the C-terminus of either full-length RNase E or RNase EΔIDR were examined by phase contrast and fluorescence (YFP) microscopy. Foci quantitation was performed using microbeJ, 326 cells were used for RNase E-YFP, and 295 cells were used for RNase EΔIDR-YFP. p-values were calculated with t-test (two-tailed, unequal variance). Error bars represent standard errors. **B.)** Representative fusion of two RNase E-YFP foci suggests phase separation. Two RNase E-YFP foci were imaged using time-lapse microscopy. Droplets were observed to fuse together and relax into a spherical droplet over 22′. >5 fusion events were observed during this time. Scale bar is 5 μm.

BR-bodies facilitate rapid mRNA turnover by compartmentalizing the RNA degradosome together with its substrate mRNAs, increasing their local concentrations (2, 4, 10). Consistent with a role in stimulating mRNA decay, RNase EΔIDR mutants have been shown to have slow mRNA decay in different bacterial species, including *E. coli* and *C. crescentus* (10, 32). To test whether the *S. meliloti* BR-body deficient mutant (RNase EΔIDR) also exhibits a slow-down in mRNA decay, we performed Rif-seq experiments to measure global mRNA half-lives (10, 33) (Fig 3A,B). In this assay, wild-type (termed RNase E henceforth) and RNase EΔIDR cells were incubated with rifampicin to block transcription, and RNA abundance was measured prior to and 1, 3, and 9 minutes after addition of rifampicin. The half-lives of RNAs can then be calculated in a genome-wide manner (10, 33). Importantly, each time-course was collected in duplicate, and tRNA is not removed from the RNA samples, as this stable RNA acts as a loading control. The bulk mRNA half-lives can be calculated from the mRNA fraction, or all the summed mRNA reads divided by total reads (mRNA+tRNA) at each time point. The mRNA fraction of the samples across the time course is then fit to a half-life equation to calculate the bulk mRNA half-lives. The bulk mRNA half-lives show that mRNA is significantly stabilized in the RNase EΔIDR mutant (from half-life of 11.7 minutes in RNase E to 25.4 minutes in the RNase EΔIDR mutant) (Fig 3B). By using the Rifcorrect software package (33), we calculated the half-lives of individual mRNAs expressed in mid-log cells (Fig 3C). Rifcorrect yielded a total of 665 mRNA half-lives that were measured in all replicates of both the RNase E and RNase EΔIDR Rif-seq datasets (Fig 3C). Of these 665 mRNAs with calculated half-lives in the RNase E and RNase EΔIDR mutant, 199/665 mRNAs were significantly more stable in the RNase EΔIDR mutant compared to RNase E (p<0.05, one-tailed t-test, unequal variance), while none of the mRNA half-lives were significantly destabilized in the RNase EΔIDR mutant. Taken altogether, the BR-body deficient mutant (RNase EΔIDR) shows a significant global slowdown in mRNA decay, suggesting that *S. meliloti* BR-bodies also promote faster mRNA degradation rates.

**Figure 3:**
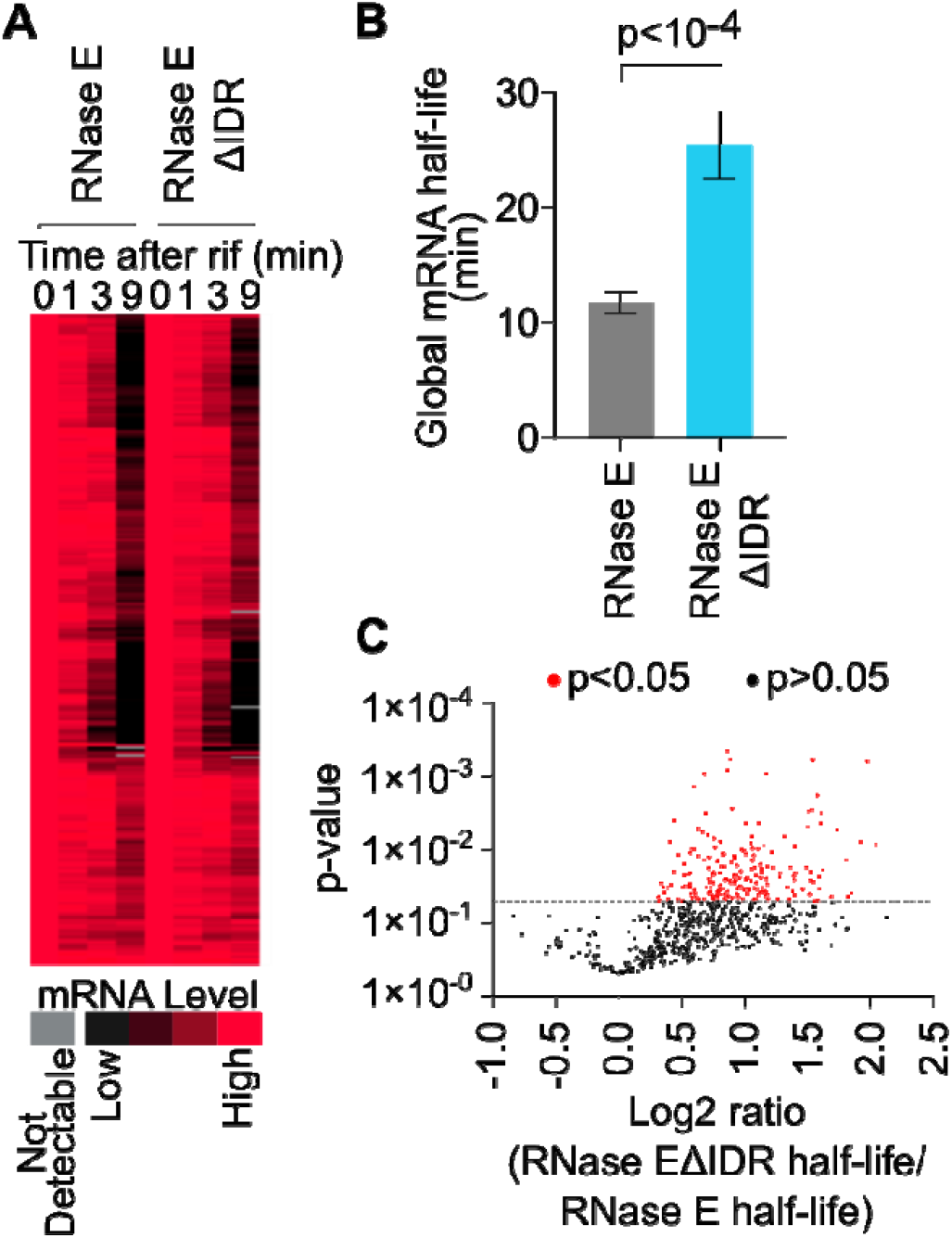
The BR-body deficient mutant (RNase EΔIDR) shows a global slowdown in mRNA decay. **A.)** Rif-seq data were collected in RNase E and RNase EΔIDR strains. RNA was extracted from cells incubated with rifampicin at the indicated time points. RNA abundance was normalized to the level in the cells prior to rifampicin treatment and displayed as a heat map by kmeans clustering. Each horizontal line represents a single mRNA. **B.)** BR-body deficient mutant (RNase EΔIDR) has a significantly higher global mRNA half-life compared to wild type (RNase E). The global mRNA half-life was calculated from a summation of mRNA reads divided by total tRNA reads (mRNA read counts + tRNA read counts) across the time points. Half-lives were estimated by curve fitting, and the average half-life between both biological replicates is presented. Error bars indicate standard deviations, and the p-value was calculated from two-tailed t-test with unequal variance. **C.)** Fold-changes of mRNA half-lives show broad stabilization in the mutant. The half-lives of 665 mRNAs could be measured in both the RNase E and RNase EΔIDR strains. 199 had significantly longer half-lives in the RNase EΔIDR compared to RNase E (p< 0.05, based on one-tailed t-test with unequal variance). Another 422 out of 665 also exhibited longer half-lives in the RNase EΔIDR, but they were not significantly different from the corresponding values in the wild type (p > 0.05). The remaining 44 mRNAs showed shorter half-lives in the mutant compared to RNase E wild type, but none were statistically significant.

### The BR-body deficient mutant (RNase EΔIDR) is sensitive to stress and defective in host colonization

*C. crescentus* BR-bodies were found to be present during normal growth but more strongly induced under different cellular stresses (4). In addition, *C. crescentus* BR-bodies promote a fitness advantage under stress-inducing conditions (4). To test the effects of stress on *S. meliloti*, we first assessed whether *S. meliloti* BR-bodies were induced under stress conditions. Similar to *C. crescentus*, we observed an increase in *S. meliloti* BR-bodies with various stresses tested [Fig S3]. The induction of BR-bodies under various stresses suggests they may also play a role in *S. meliloti* fitness. Therefore, we compared the growth of wild type (RNase E) and the BR-body deficient mutant (RNase EΔIDR) under various stresses (Fig 4). To ensure that observed phenotypes were due to the deletion of the RNase E IDR, and not a secondary mutation, we repaired the IDR in the BR-body deficient mutant (RNase EΔIDR) background, generating the *rne*^+^ strain. Under permissive conditions at 28°C, we did not observe any significant differences in viability or colony size on solid media (Fig 4) or growth rates in liquid media (Fig S1). Upon incubating the cells with a variety of stresses, we observed with the mutant a reduction in colony size when grown at higher temperature (37°C), and strong reduction in CFUs when grown in the presence of ethanol (Fig 4). In all cases, the *rne*^+^ strain grew the same as the RNase E wild type, indicating that the IDR deletion causes the stress tolerance defect. This suggests that *S. meliloti* BR-bodies provide a fitness advantage under various stresses and that the stress tolerance phenotype observed in *C. crescentus* is likely conserved across bacteria.

**Figure 4:**
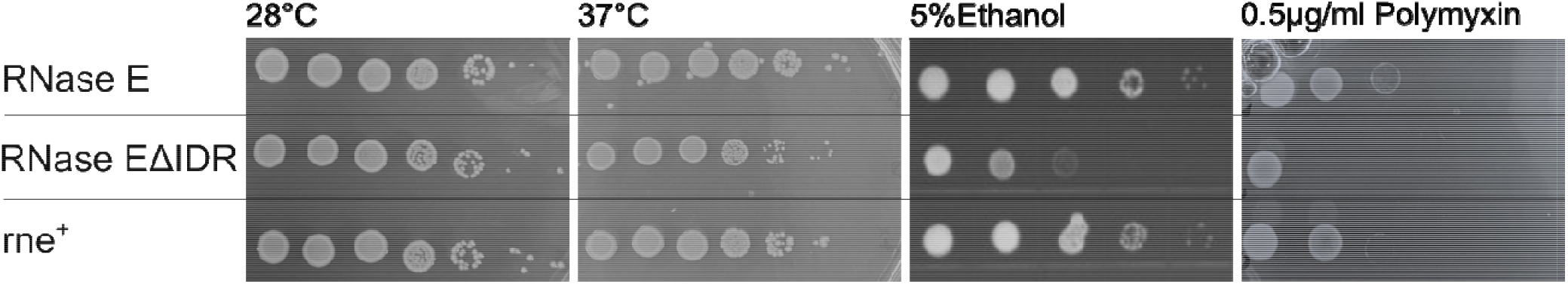
The BR-body deficient mutant (RNase EΔIDR) is sensitive to multiple stresses. The BR-body deficient (RNase EΔIDR) mutant is more sensitive to various stresses compared to RNase E and *rne*^+^ (derivative of RNase EΔIDR in which the IDR deletion was repaired at the *rne* locus).

While *C. crescentus* is a free-living bacterium, *S. meliloti* is a facultative endosymbiont that can invade root tissue and colonize nodules, fixing nitrogen for the host plant to promote growth. To achieve effective symbiosis, *S. meliloti* must adapt to and survive host defenses. The model host *M. truncatula* is known to produce several cationic nodule-specific cysteine-rich□(NCR) peptides as a part of its innate immune response against bacteria (21–24). We used polymyxin B, a widely available cationic peptide antibiotic to mimic these NCR peptides and found that the BR-body deficient mutant (RNase EΔIDR) was sensitive to polymyxin (Fig 4), indicating that BR-bodies may promote a fitness advantage during host colonization. To study the role of BR-bodies during infection, *M. truncatula* seedlings were inoculated with wild type (RNase E), BR-body deficient mutant (RNase EΔIDR), or *rne*^+^ *S. meliloti* cells, and the plants were grown and monitored on nitrogen-free medium for six weeks (Fig 5). Seedlings inoculated with the BR-body deficient mutant (RNase EΔIDR) grew poorly compared to RNase E, suggesting that the mutant is unable to form effective symbiosis (Fig 5A). The poor plant growth observed with the BR-body deficient mutant (RNase EΔIDR) was fully alleviated with the *rne+* strain, demonstrating that the symbiosis defect was caused by the deletion of RNase E’s IDR (Fig 5A). Furthermore, while we observed similar numbers of root nodules in plants inoculated with different strains (Fig 5B), plants inoculated with the BR-body deficient mutant (RNase EΔIDR) exhibited a higher proportion of white nodules instead of healthy pink nodules, suggesting a failure to fix nitrogen (Fig 5C). To examine the symbiosis defect more closely, we performed competitive colonization experiments to compare the relative fitness of the wild-type and RNase EΔIDR strains. We generated 1:1 mixtures of wild type and the BR-body deficient mutant (RNase EΔIDR), inoculated plants with the mixtures, and recovered bacteria from individual nodules 28 days later (Fig 5D). Each strain carried a unique antibiotic resistance marker for rapid phenotypic identification. As a control, we also competed two distinctly marked wild-type RNase E strains against each other. We found that the BR-body deficient mutant (RNase EΔIDR) showed a large reduction in root nodule occupancy compared to RNase E. In contrast, equal ratios of wild-type strains were recovered in competitions between two with wild-type RNase E (Fig 5D, Fig S4), suggesting that the drug markers did not lead to differences in fitness during symbiosis. Thus, the reduction in nodule occupancy by the BR-body deficient mutant (RNase EΔIDR) compared to wild type suggests that BR-bodies promote fitness during host colonization (Fig 5D, Fig S4).

**Figure 5:**
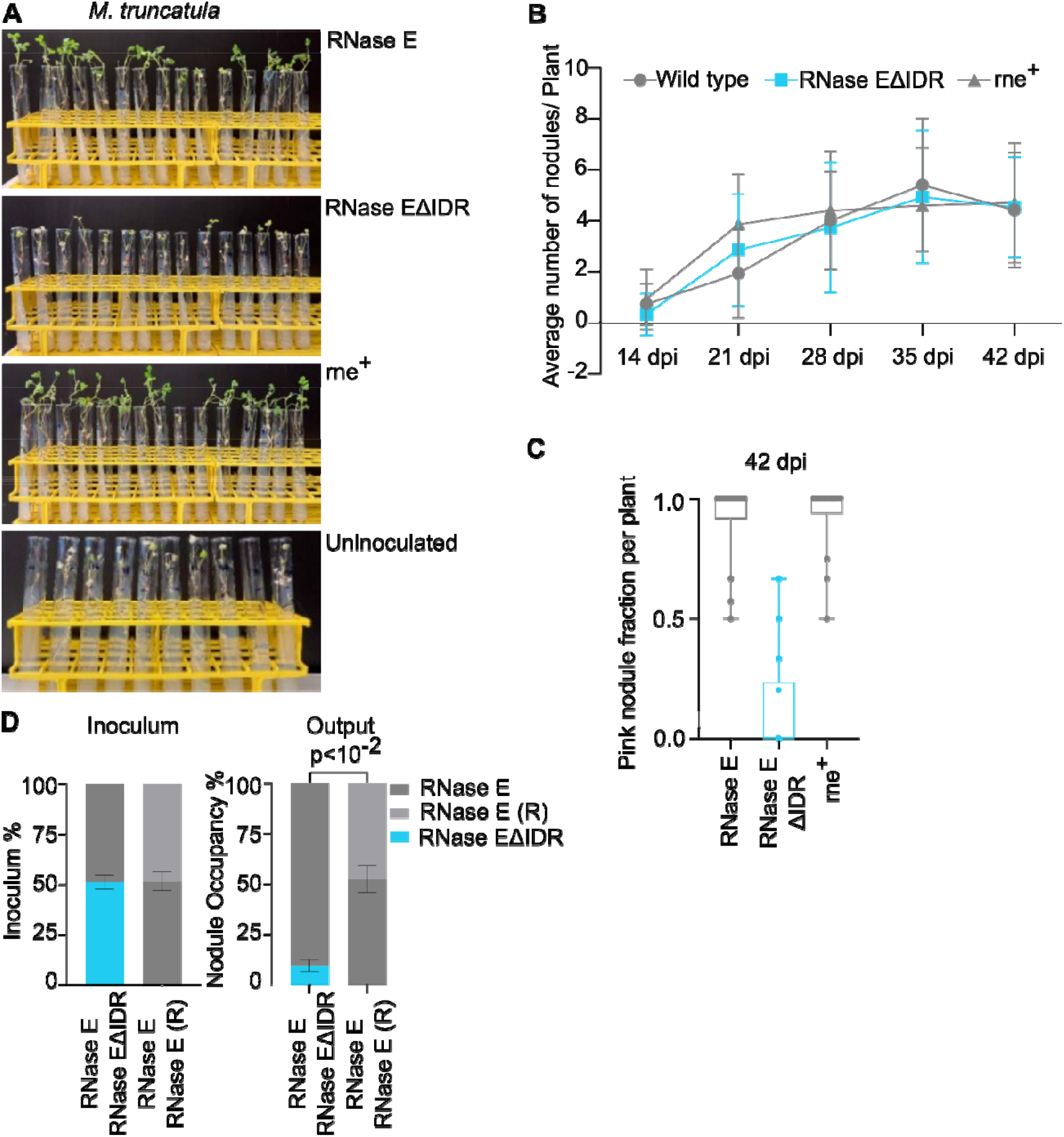
The BR-body deficient mutant (RNase EΔIDR) is defective in *M. truncatula* root colonization and symbiosis. **A.)** *M. truncatula* plants were photographed after being inoculated with the indicated strain of *S. meliloti* and grown on nitrogen-free media for six weeks. **B.)** Average number of root nodules on *M. truncatula* inoculated with *S. meliloti* strains. The number of root nodules per plant and their colors were recorded every seven days, starting at 14 days post-infection. Error bars represent standard deviations. Uninoculated plants did not develop any nodules. **C.)** Fraction of pink nodules per plant inoculated with each strain 42 days post-infection. Seedlings inoculated with strains expressing wild-type RNase E had significantly higher fractions of pink nodules compared to seedlings inoculated with RNase EΔIDR strains; p < 1x10^−4^ (two tailed, Mann-Whitney test). **D.)** RNase EΔIDR is less competitive in colonizing plant root nodules. A competitive symbiosis assay was used to compare the relative colonization efficiency of wild type (RNase E) and the BR-body deficient mutant (RNase EΔIDR). The right plot shows mean percentages (± standard deviations) of root nodules colonized by each bacterial strain in distinct competitions, measured 28 days after seedlings were inoculated with equal mixtures of two strains. RNase E (R) indicates that strains marked with resistance to spectinomycin or neomycin were used in the competition. Ratios in the left plot were derived from colony counts when the inoculating mixtures were plated. Relative nodule occupancy when RNase E competed against RNase EΔIDR was significantly different than that when RNase E competed against RNase E (p< 0.01, based on two-tailed t-test with unequal variances). A total of three competition replicates were used for wild type vs. RNase EΔIDR (n=318 nodules) and seven competition replicates for wild type vs. wild type (n=587 nodules). As the spectinomycin and neomycin resistance markers did not affect fitness in the competitive assay (Fig S4), we combined all wild type vs. wild type or wild type vs. RNase EΔIDR competitions for data analysis.

## Discussion

### BR-body cellular functions are conserved in α-proteobacteria

While RNase E’s IDR was found to be necessary and sufficient for BR-body assembly in *C. crecscentus*, and RNase E proteins contain IDRs across bacteria (2), the IDR’s importance in BR-body assembly has not been directly tested in other species. In *S. meliloti*, we showed that the C-terminal IDR is necessary for BR-body assembly *in vivo* (Fig 2), allowing our RNase EΔIDR mutant to act as a BR-body deficient mutant (Fig 2). In addition, BR-bodies in *S. meliloti* appear to be dynamic, liquid-like condensates as we observed events of droplet fusion (Fig 2). In accordance with this, we found that BR-bodies increase mRNA decay rates across the transcriptome (Fig 3) and appear to be stimulated by mRNAs which are not being efficiently translated in polysomes (Fig S2), as observed in *C. crescentus* (4, 10). BR-bodies are constitutively present during exponential growth; however, they are also stimulated in the presence of cell stresses, whereby they promote stress resistance (Fig 4) (4). This suggests that despite the diverse sequences and lengths of the IDRs across species (2), the molecular and cellular functions of BR-bodies appear to be similar across species.

### BR-bodies promote plant colonization and symbiosis

While RNase E mutants with ΔIDR or Tn-insertions that truncate the IDR have been found to have reduced fitness during host colonization in multiple species (12, 14–18), there has not been direct demonstration that these mutations prevented BR-body formation. Therefore, this study provides the first evidence that BR-body function can promote fitness during host colonization (Fig 6). While the *S. meliloti* BR-body deficient RNase EΔIDR mutant can colonize *M. truncatula* roots, it was readily outcompeted by the RNase E wild type in a direct competition assay, suggesting there is a significant loss in fitness during symbiosis (Fig 5). While the RNase EΔIDR mutant appears to have similar growth to wild type (RNase E) in unstressed conditions *in vitro*, we hypothesize that the stress sensitivity makes the bacterial cells more susceptible to the plant’s innate immune defenses, such as antimicrobial peptides (24). In addition, the *S. meliloti* BR-body deficient mutant (RNase EΔIDR) failed to stimulate plant growth and appeared to generate white root nodules, suggesting a defect in nitrogen fixation (Fig 5). Previous studies demonstrated that sRNAs and their chaperone Hfq play important roles in *S. meliloti* during symbiosis by performing post-transcriptional regulation on key mRNAs (26, 27, 34). BR-bodies may be important organizers of sRNA regulation, as sRNAs and Hfq were found to be enriched in *C. crescentus* BR-bodies (10, 11). In addition, RNase E’s IDR is required for sRNA regulation in multiple *E. coli* sRNAs (35, 36). Interestingly, a similar RNase E IDR truncation mutant in *Brucella abortus* showed attenuated mouse infection and altered sRNA levels (12), suggesting BR-bodies and sRNAs may also coordinate activities in animal pathogens.

**Figure 6:**
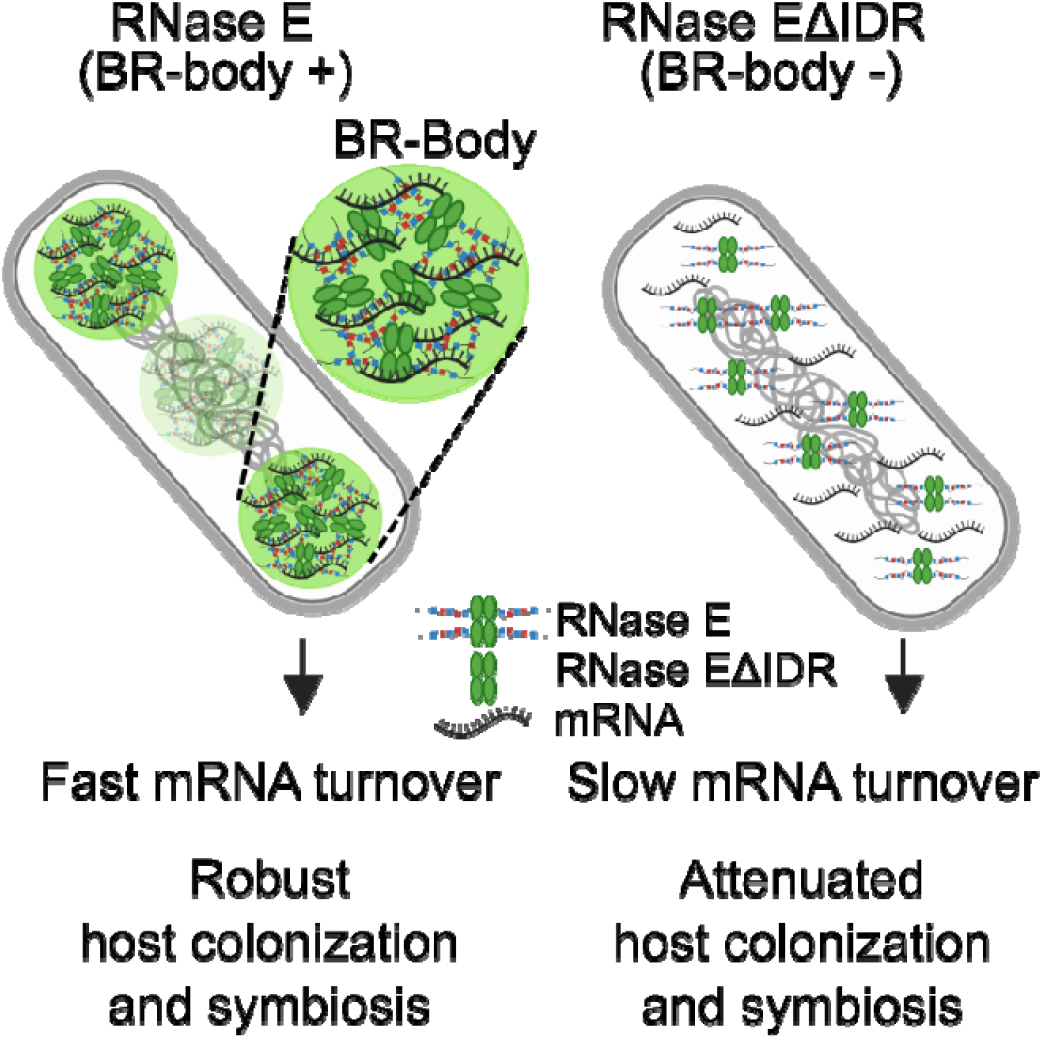
BR-bodies stimulate mRNA decay and promote host colonization. A summary of molecular and cellular phenotypes impacted by the BR-body deficiency, as depicted in both wild type and the RNase EΔIDR mutant.

## Materials and Methods

### Sinorhizobium meliloti cell growth

All *S. meliloti* strains used in this study (Table 1) were derived from Rm1021 (wild-type RNase E) and were grown at 28°C in Luria-Bertani broth (LB), peptone-yeast extract (PYE), or tryptone-yeast extract (TY) medium, supplemented with gentamicin (Gent), neomycin (Nm), spectinomycin (Sp), and/or streptomycin (Strep) when appropriate (37). All plasmids used in this study are in Table 2. Optical density (OD) of liquid cultures was measured at 600 nm in a cuvette using a Nanodrop 2000C spectrophotometer. Strains were analyzed at mid-exponential phase of growth (OD of 0.3–0.6).

**Table 1.** *Sinorhizobium meliloti* strains used in this study

| Strains | Relevant genetic markers, features, and/or description | Construction, source, or reference<sup>a</sup> |
| --- | --- | --- |
| Rm1021 | SU47 derivative, Sm<sup>R</sup> (progenitor of strains listed below) | (47) |
| JS9 | *rne::pRNE(Sm)-YFPC-r (rne-yfp)* | (4) |
| JS377 | *rne::pJSE2 [rne(NTD)-yfp]* | pJSE2 mated into Rm1021 |
| JOE2612 | *podJ2*(SMc02231)..$\Omega$ (Sp<sup>R</sup> wild-type strain) | (41) |
| JOE6312 | *podJ2*(SMc02231)..*nptII* (Nm<sup>R</sup> wild-type strain) | Allelic replacement in Rm1021 using pJC739 |
| JOE6516 | *rne*ΔCTD | Allelic replacement in Rm1021 using pJSE3 |
| JOE6526 | *podJ2*(SMc02231)..*nptII rne*ΔCTD | Allelic replacement in JOE6312 using pJSE3 |
| JOE6550 | *podJ2*(SMc02231)..$\Omega$ *rne*ΔCTD | JOE6516 x $\Phi$(JOE2612), select for Sp<sup>R</sup> |
| JOE6825 | *podJ2*(SMc02231)..*nptII rne*<sup>+</sup> | Allelic replacement in JOE6256 using pJSE4 |
| JOE6826 | *podJ2*(SMc02231)..$\Omega$ *rne*<sup>+</sup> | Allelic replacement in JOE6550 using pJSE4 |
| JOE6831 | *rne*<sup>+</sup> | Allelic replacement in JOE6516 using pJSE4 |

<sup>a</sup> $\Phi$ indicates generalized transduction, as mediated by bacteriophage $\Phi$ N3. For example, JOE6516 x $\Phi$ (JOE2612) means that a bacteriophage lysate made from JOE2612 was used to infect JOE6516.

**Table 2.** Plasmids used in this study

| Plasmid | Relevant genetic markers, features, and/or description | Construction, source, or reference<sup>a</sup> |
| --- | --- | --- |
| pJQ200sk | Counter-selectable vector for allelic replacement, *sacB*, Gm<sup>R</sup> | (40) |
| pVYFPC-4 | Integrating plasmid with multiple cloning site under control of the vanillate promoter, Gm<sup>R</sup> | (39) |
| pRNE(Sm)-YFPC-4 | Integrating plasmid carrying the last 1101 bp of *rne* from *S. meliloti* Rm1021 in frame with a C-terminal YFP tag, derived from pYFPC-4 | (4) |
| pJC739 | pJQ200sk-*podJ2*..*nptII*, for inserting *nptII* (Nm/Km resistance marker) after *podJ2* gene | This study |
| pJSE2 | Integrating plasmid carrying the last 512 bp of *rne* NTD from *S. meliloti* Rm1021, for fusing the N-terminal domain of RNase E in frame with a C-terminal YFP tag, derived from pYFPC-4 | This study |
| pJSE3 | pJQ200sk-*rne*ΔCTD, for allelic replacement to delete C-terminal domain of RNase E | This study |
| pJSE4 | pJQ200sk-*rne*<sup>+</sup>, for allelic replacement to repair mutation in *rne* | This study |

<sup>a</sup> Plasmid sequences are available in GenBank format in Supplemental Information.

For doubling time calculations Rm1021 derivatives were grown as 2 mL cultures in TY media. Culture tubes were incubated in a shaker incubator at 240 rpm at 28°C. Absorbance at 600 nm was measured every 1 hour using a Thermo Scientific Genesys 20 spectrophotometer. Averages and standard deviations were calculated from readings of three biological replicates of each genotype (RNase E, RNase EΔIDR or *rne*^+^).

For acute drug treatment, the log phase cells were treated with rifampicin (100 μg/mL) or chloramphenicol (200 μg/mL) for 30′. For serial dilution assays, including cell stress assays, cells were grown overnight in TY media and diluted to OD of 0.5, followed by ten-fold serial dilutions. 5 μL each of select dilutions were spotted on plates with the indicated composition and incubated at 28 or 37°C.

### Strain and plasmid construction

#### JS9: Rm1021 *rne::rne-yfp* (Gent^R^)

This strain was initially generated in a previous study by Al-Husini et al, 2018 (4).

#### JS377: Rm1021 *rne::(rne(NTD)-YFP* (Gent^R^)

*S. meliloti* RNase E NTD-YFP fusion (RNase EΔIDR-YFP) was generated by amplifying the last 500 bp encoding the RNase E N-terminal domain (NTD) using the following primers designed using the j5 DNA assembly design automation software (38):

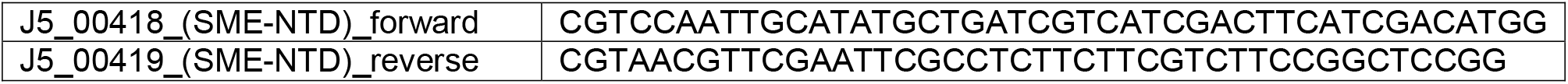

The resulting PCR product and pYFPC-4 plasmid (39) were both cut with NdeI and EcoRI, ligated, and transformed into *E. coli* DH5α (Invitrogen). Gent^R^ colonies were then screened by PCR for the NTD insert, and the purified plasmid was verified by Sanger sequencing (Genewiz). This pJSE2 plasmid was then mated into Rm1021 by triparental mating. Desired transconjugants were selected on Gent/Strep plates and verified by PCR and YFP imaging.

#### JOE6516: Rm1021 *rne*ΔIDR

*S. meliloti* RNase EΔIDR mutants and *rne*^+^ correction strains were generated via two-step allelic replacement by homologous recombination, using plasmids derived from pJQ200sk (40); pJSE3 was used to delete the region coding for the C-terminal IDR. pJSE3 was generated as follows.

First, 1000 bp upstream and 1000 bp downstream of the region encoding the RNase E C-terminal domain (CTD) were amplified by PCR from the Rm1021 genome using the following DNA primers designed using the j5 DNA assembly design automation software (38):

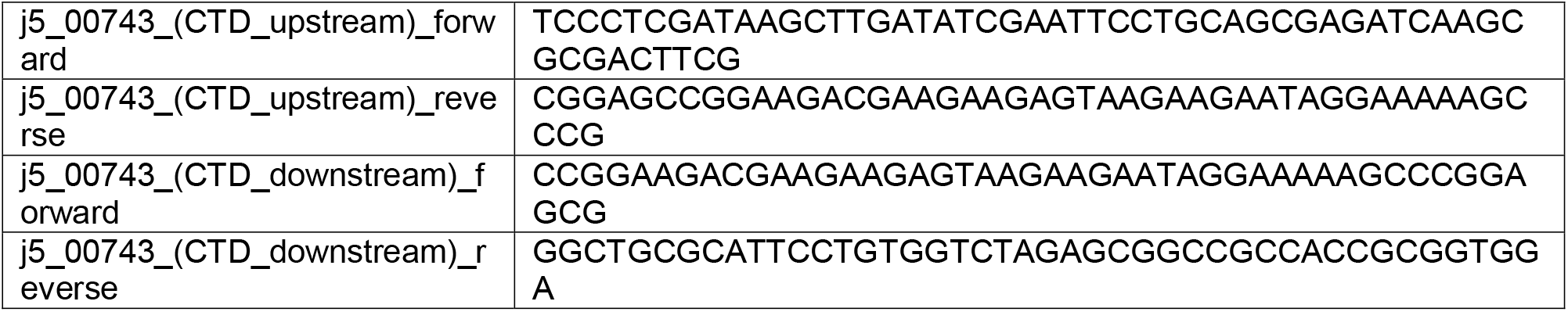

Next, the resulting upstream and downstream PCR fragments were gel purified, assembled into pJQ200sk via Gibson cloning, and transformed into *E. coli* DH5α (Invitrogen). Gent^R^ colonies were then screened by PCR for both inserts, and purified plasmids were verified by Sanger sequencing (Genewiz).

JOE6526 (Rm1021 *podJ2*..*nptII rne*ΔIDR) was also constructed by using pJSE3 to generate the deletion in JOE6312 (Rm1021 *podJ2*..*nptII*), while JOE6550 (Rm1021 *podJ2*..Ω *rne*ΔIDR) was constructed by transducing the *podJ2*..Ω allele from JOE2612 (41) into JOE6516.

JOE6312 was generated by introducing the *nptII* gene into Rm1021, downstream of SMc02231, using the pJQ200sk-derived plasmid pJC739, which was constructed by amplifying the *nptII* gene from pBbB2k-GFP (42) with primers nptII -136F and nptII +3R (see sequences below), digesting the resulting PCR product with BamHI, and inserting the fragment into the BamHI site of pJC377 (parent of pJC382) (41).

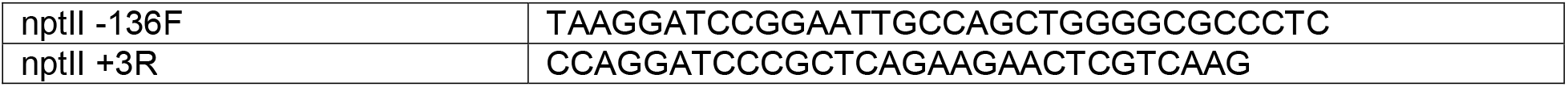

#### JOE6831: Rm1021 *rne*^*+*^

Plasmid pJSE4 was used to reintroduce the wild-type *rne* allele into JOE6516. pJSE4 was generated by amplifying from 1000 bp upstream to 1000 bp downstream of the region encoding the RNase E CTD from the Rm1021 genome using the following DNA primers designed using the j5 DNA assembly design automation software (38):

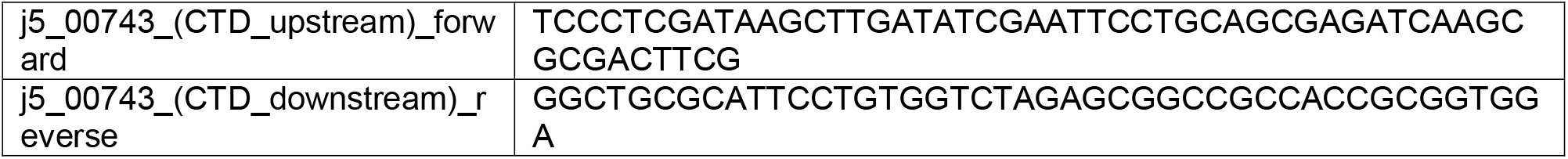

Next, the resulting PCR product was gel purified, inserted into pJQ200sk via Gibson assembly, and transformed into *E. coli* DH5α (Invitrogen). Gent^R^ colonies were then screened by PCR for the full insert, and purified plasmids were verified by Sanger sequencing (Genewiz).

JOE6825 (Rm1021 *podJ2*..*nptII rne*^*+*^), and JOE6826 (Rm1021 *podJ2*..Ω *rne*^*+*^) were similarly constructed via two-step allelic replacement from JOE6526 or JOE6550 (Rm1021 *podJ2*..Ω *rne*ΔIDR), respectively.

### Cell imaging and analysis

For imaging, cells were immobilized on 1.5% agarose pads made with 1x M9 salts on microscope slides, and images were collected using a Nikon Eclipse NI-E with CoolSNAP MYO-CCD camera and 100x Oil CFI Plan Fluor (Nikon) objective, driven by NIS-Elements software. The Chroma 96363 filter set was used for YFP. For time lapse experiments, images were taken at 1′ intervals with manual focus over a maximum time period of 30′ and approximately 5-8 fusion events were observed per movie.

Cell image analysis was performed using microbeJ (43). For automated foci detection, the maxima foci function of microbeJ was used. The tolerance and Z-score settings were manually adjusted to identify and outline foci on a test image then removed aberrant foci with area <0.01 μm^2^ and length >1 μm. The segmentation option was also used to split adjoined foci. For a given set of images, the same tolerance/Z-score parameters were applied to quantify the number of foci per cell with a minimum of 50 cells.

### RNA decay rates by RNA-seq

Cells were grown in TY media overnight. The next day, mid-log phase cultures (OD600 0.3 - 0.5) were treated with 10 mg/mL rifampicin to achieve a final concentration of 200 μg/mL. Before adding rifampicin, 1 mL of culture (0′) was removed and mixed with 2 mL of RNAprotect Bacterial Reagent (QIAGEN), vortexed, and incubated at room temperature for 5′. This step was repeated at 1′, 3′, and 9′ after rifampicin was added. The cells were pelleted at (20,000xg) for 1′ and resuspended in 1 mL of 65°C TRizol Reagent (Ambion) and incubated at 65°C for 10′ in a heat block. 200 μL of chloroform was added to the samples, and the tubes were incubated at room temperature for 5′ before spinning at 20,000xg for 10′. RNA samples were chloroform extracted once and precipitated using isopropanol (1x volume isopropanol, 0.1X volume 5 M sodium acetate, pH 5.2) overnight at −80°C. The RNA samples were spun at 20,000 x g at 4°C for 1 hour; pellets were washed with 80% ethanol for 10′, air dried, and resuspended in 10 mM Tris-HCl (pH 7.0). The RNA-Seq libraries were made using 5 μg of total RNA samples, and library construction was performed according to published protocol (44). Raw sequencing data is available in the NCBI GEO database with accession number GSE251703. To measure RNA-decay rates, the fraction of mRNA remaining was calculated as the RPKM of each time point divided by the RPKM measured in the untreated 0′ sample (Table S1). For bulk mRNA half-life calculations of RNA-seq data the fraction of all mRNA reads was compared to the total fraction of reads which includes a majority of stable tRNA reads (Table S2).

### Symbiosis assays with *M. truncatula*

*M. truncatula* seedlings were inoculated with Rm1021 or derivatives and grown in nitrogen-free media as previously described (37). Root nodules were counted every week for six weeks, starting at two weeks post-inoculation. Photographs of the plants were captured at the end of six weeks. Competitive colonization assays were also performed as previously described (37). Briefly, equal volumes of two cell suspensions (*S. meliloti* Rm1021 and its derivatives) were mixed for inoculating *M. truncatula* seedlings. Strains were marked with resistance to spectinomycin or neomycin for phenotypic identification. Bacteria were recovered from nodules 28 days post-inoculation to determine the dominant strain in each nodule (>60 nodules per trial, 3 or more trials per competition) (Table S3). For one of the RNase E vs. RNase EΔIDR competitions (JOE2612 vs. JOE6550), both strains carried the spectinomycin resistance marker. In that competition, the genotype of individual isolates from the inoculating mixture and the nodule extracts was determined by PCR using the primers SMc01336 1825F and SMc01336 +285R (sequences shown below). Wild-type *rne* yielded a 1235-bp product, while the ΔIDR mutation yielded a 458-bp product.

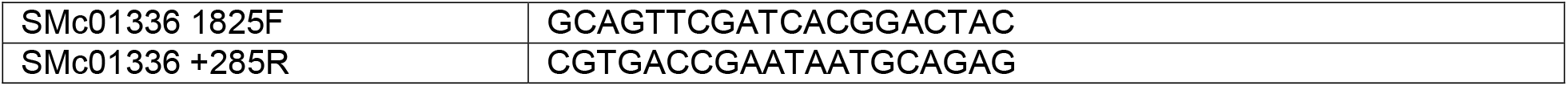

All the information on statistical analysis can be found in Table S4.

## Author contributions

JMS and JCC designed the study, obtained funding. KSM and AG generated plasmids. KSM, AG, NA-H and JMS performed imaging experiments. KSM and IWR generated Rif-seq libraries. KSM and JMS analyzed Rif-seq data. KSM, ZH, RAC, NS-R, LHM, EN and JCC performed stress experiments. JLH, SM, RAC, ZH and JCC performed plant colonization experiments. KSM, JCC and JMS analyzed data. KSM, JMS and JCC wrote paper.

## Acknowledgements

The authors acknowledge support from NIH funding GM124733 to JMS and SC3-GM096943 to JCC. RAC received support from grants T34-GM008574, T34-GM145400, and R25-GM050078. NS-R also received funding from grant T34-GM008574. Rowan Poppen-Eagan and Sam Cuevas helped perform initial characterization of mutant strains.

## Notes

### Competing Interest Statement

The authors have declared no competing interest.

