## Supplemental figures for "*Sinorhizobium meliloti* BR-bodies promote fitness during host colonization"

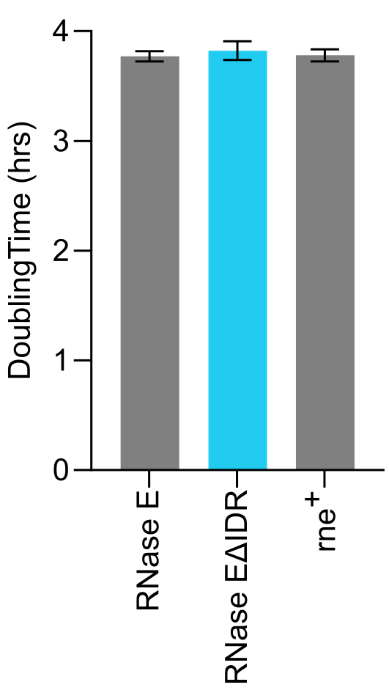

**Figure S1: Growth rates for *S. meliloti* strains in TY media.** The growth rates of two-mL cultures were determined by measuring the optical density at 600 nm (OD600) every hour, starting at 0.05. Exponential curve fits were performed in mid-log phase (OD600 0.275-0.550). The average doubling time was calculated, and the error bars represent standard deviations from three replicate growth curves.

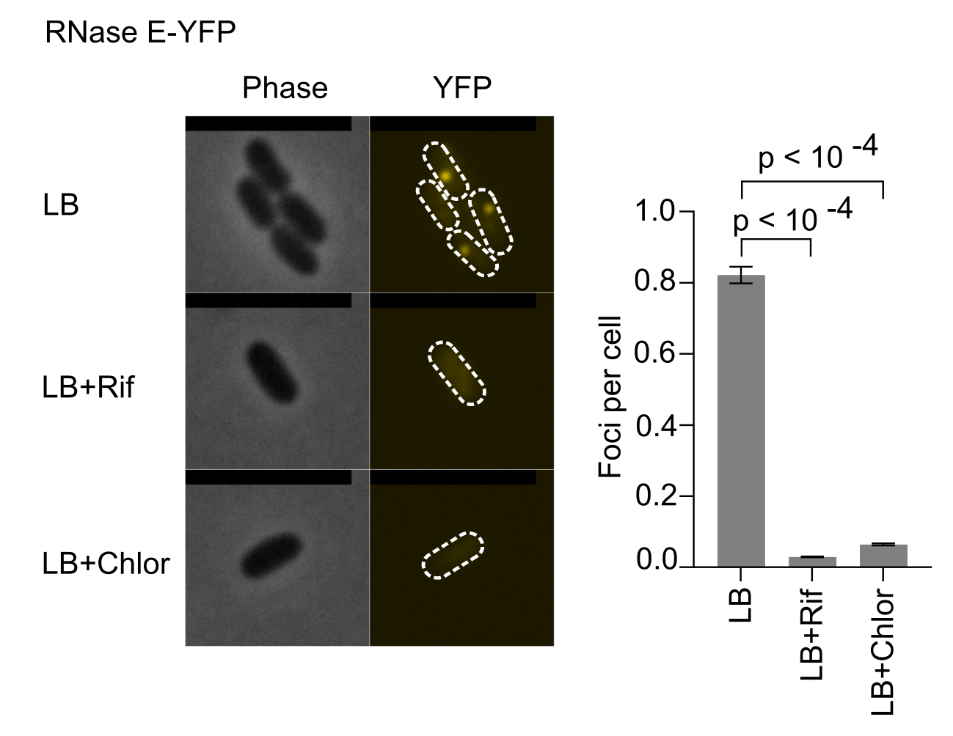

**Figure S2:** ***S. meliloti* BR-bodies are mRNA-dependent.** Cells expressing RNase E-YFP and grown in LB were treated with 100 µg/mL of rifampicin (Rif) for 30′ to deplete mRNA or 200 µg/mL of chloramphenicol (Chlor) for 30′ to arrest translation and accumulate mRNA in polysomes, prior to examination by microscopy and detection of fluorescence foci. Foci quantitation was performed using microbeJ to compare treated and untreated cells. 1240 cells were used for untreated cells, 482 cells were used for rifampicin-treated cells, and 467 cells were used for chloramphenicol-treated cells. p-values were calculated with t-test (two-tailed, unequal variance). Error bars represent standard errors. Scale bar is 5 μm.

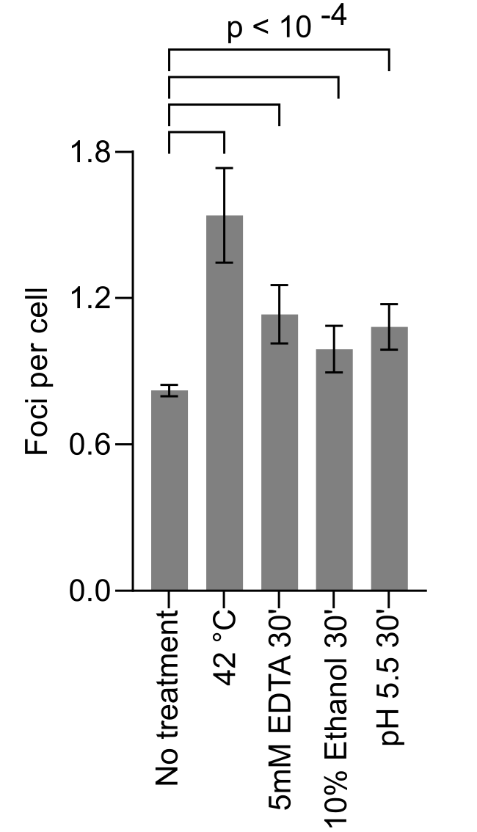

**Figure S3: *S. meliloti* BR-bodies can be strongly induced under stressful conditions.** RNase E-YFP cells were grown in TY media and then treated with the indicated stresses for 30′ before being placed on TY 1.5% agarose pads for imaging. >100 cells were analyzed for each condition, and the average foci/cell was calculated using microbeJ. The error bars represent standard errors. p-values were calculated with t-test (one-tailed, unequal variance).

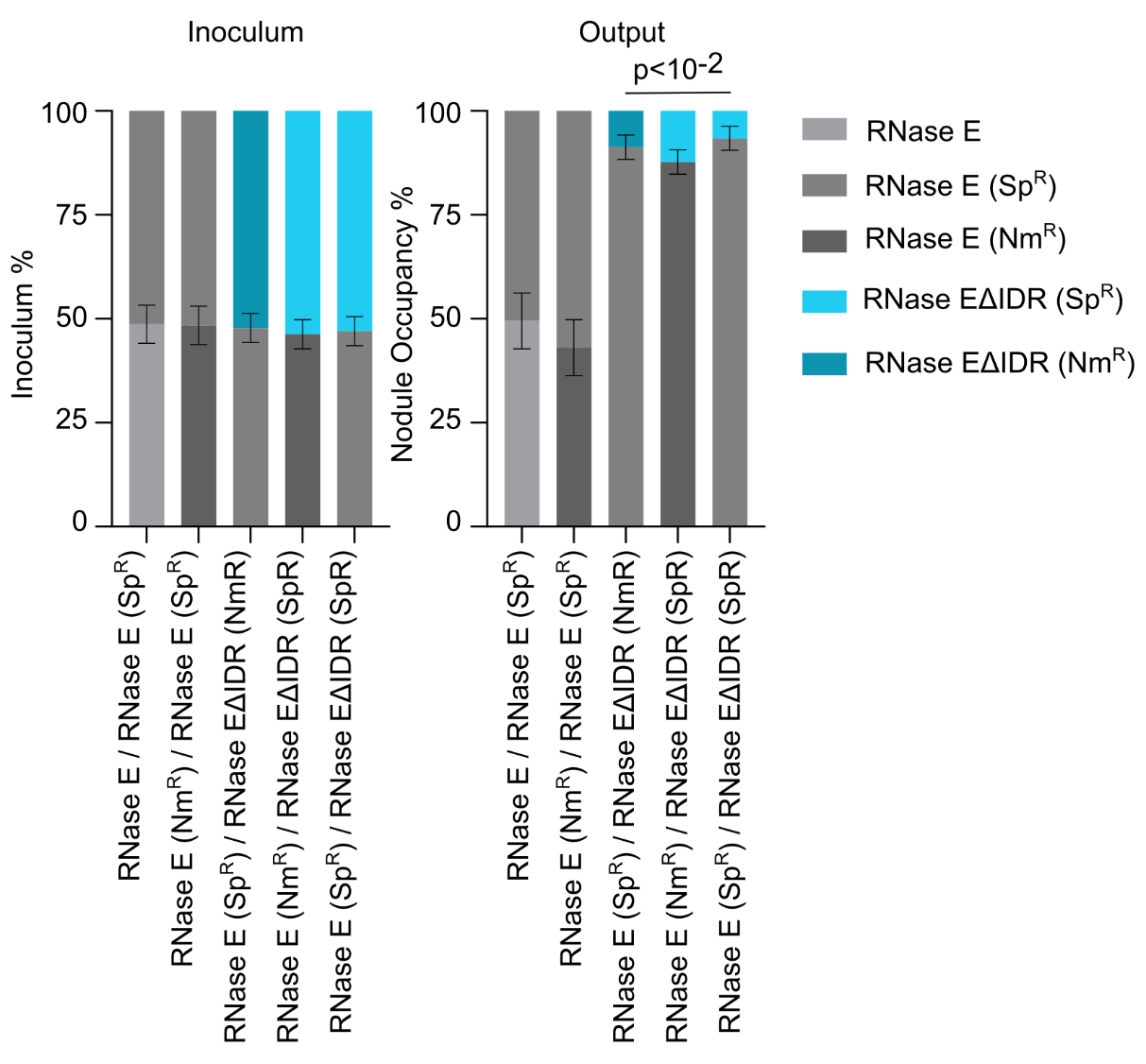

**Figure S4: RNase EΔIDR is less competitive in colonizing plant root nodules.** A competitive symbiosis assay was used to test the relative colonization of wild type (RNase E) and the BR-body deficient mutant (RNase EΔIDR). Proportions of *M. truncatula* root nodules colonized by each bacterial strain were measured after seedlings were inoculated with equal mixtures of two strains. RNase E and RNase EΔIDR strains marked with resistance to spectinomycin or neomycin were used in the competition. Percentages of strains in each inoculum were derived from colony counts when the inoculating mixtures were plated on selective media. Error bars represent standard deviations.
